# Mesenchymal stem cell derived exosomes enhance lymphangiogenesis via exosomal transfer of Ang-2/Tie2

**DOI:** 10.1101/466987

**Authors:** Ting Zhao, Zhixin Yan, Jinwen Liu, Hui Sun, Yifei Chen, Yan Tao, Wenrong Xu, Hui Qian, Yongmin Yan

## Abstract

Mesenchymal stem cell derived exosomes (MSC-Ex) are nanosized membrane-bound extracellular vesicles found in MSC conditioned medium, that have yielded beneficial effects in several experimental models of organ injury. However, the therapeutic value and mechanism of MSC-Ex in lymphedema is poorly understood. Here we find that human umbilical cord MSCs derived exosomes (hucMSC-Ex) treatment contributed to the regeneration of LYVE-1 positive lymphatic vessels and reduction of lymphedema in a mouse model of tail lymphedema. Following uptake, exosomal lymphangiogenic factors (angiopoietin (Ang)-2 and Tie2) are taken up by HDLECs and promoted HDLECs proliferation, migration, and tube formation in vitro. We also find that exosomal Ang-2 and Tie2 exert a prolymphangiogenic effect on HDLECs through upregulating Prox1 and VEGFR3/p-Akt expression. In conclusion, our result unravel a previously unappreciated prolymphangiogenic role of hucMSC-Ex in lymphedema therapy and provide a new mechanism of Ang-2 in therapeutic lymphangiogenesis.

## Introduction

Lymphedema is seen as pathologic swelling after localized lymphatic fluid retention that results from lymphatic system damage, and it affects approximately 250 million people worldwide.(Greene, Voss et al., 2017, Schulze, Nacke et al., 2018) Although nonoperative and operative treatments for lymphedema are well established, these procedures typically provide only partial relief from limb swelling, functional impairment, and the risk of cellulitis.(Schaverien & Aldrich, 2018) At present, no definitive treatments are available for lymphedema.(Stuiver, Ten Tusscher et al., 2017) Thus, new therapies to treat chronic lymphedema are urgently needed.

Recently, preclinical animal models and clinical trials have shown that cell-based therapy using mesenchymal stem cells (MSCs) is a promising new therapeutic approach for treating lymphedema because these cells have multiple differentiation potentials and capacity for growth factor secretion.(Conrad, Niess et al., 2009, Qi & Pan, 2015, Zhou, Wang et al., 2017) Although two human studies have investigated the possible beneficial effects of stem cells on lymphedema,(Hou, Wu et al., 2008, Maldonado, Perez et al., 2011) malignant transformation is still a concern.(Jeong, Han et al., 2011, Zhou, Bosch-Marce et al., 2006) MSC-derived exosomes (MSC-Ex) are nanosized membrane-bound extracellular vesicles found in MSC conditioned medium, that have yielded beneficial effects in several experimental models of organ injury.(Sun, Shi et al., 2018, Willis, Fernandez-Gonzalez et al., 2018, Xiao, Wang et al., 2018, Yeo, Lai et al., 2013) In comparison with MSC transplantation, MSC-Ex therapies are characterized by fewer immune responses, increased safety, and ease of storage, shipment, and administration.(Lai, Chen et al., 2011) We also demonstrated that MSC-Ex could alleviate liver and renal injury, as well as type-2 diabetes and wound healing, and have similar protective and regenerative properties as their cellular counterparts.(Jia, Liu et al., 2018, Jiang, Tan et al., 2018, Shi, Xu et al., 2017, Sun et al., 2018, Yan, Jiang et al., 2017) Additionally, exosomes from adipose-derived stem cell (ADSCs) were reported to promote proliferation and migration of lymphatic endothelial cells (LECs) in vitro.(Wang, Wang et al., 2018) However, the effects and mechanisms of MSC-Ex on lymphedema remain largely unexplored.

MSC-Ex exert effects via delivering MSC-derived cargos (i.e., DNAs, mRNAs, miRNAs, and proteins) to host cells, which lead to changed behaviors of recipient cells to enhance tissue repair.(Liu, Lin et al., 2018, Mayourian, Ceholski et al., 2018, Valadi, Ekstrom et al., 2007, Yan et al., 2017) Thus, it will be necessary to determine the factors that control hucMSC-Ex mediated lymphangiogenesis and their antilymphedemic effects. So far antilymphedema studies have only used bone marrow MSC (BM-MSCs) and ADSCs.(Ackermann, Wettstein et al., 2015, Toyserkani, Jensen et al., 2018, Zhou et al., 2017) Human umbilical cord-derived MSCs (hucMSCs) are of interest in regenerative medicine because of their low cost, minimal invasiveness, and convenient isolation methods.(Bartolucci, Verdugo et al., 2017, Ding, Chang et al., 2015, Li, Xia et al., 2015) Thus, hucMSC-Ex may represent an ideal cell-free therapy for lymphedema. The aim of the present study was to characterize the potential of hucMSC-Ex to enhance lymphangiogenesis and reduce lymphedema in vivo, and to identify the key exosomal prolymphangiogenic factors to provide a mechanistic basis for optimizing future hucMSC-Ex-based lymphedema therapies.

## Results

### hucMSC-Ex Reduces Lymphedema In Vivo

Transmission electron microscopy (TEM) and nanoparticle tracking analysis (NTA) were utilized to confirm that the hucMSC-Ex had spheroid morphology (diameter 30-100 nm; Figure 1A and 1B). Western blot analyses confirmed the presence of exosomal markers (CD9 and CD63) in the hucMSC-Ex and their cellular counterparts (Figure 1C). Acquired lymphedema was consistently induced by microsurgical ablation of the major lymphatic conduits in the murine tail. A significant degree of edema developed within one week after the microsurgery. To evaluate the therapeutic effect of hucMSC-Ex on the lymphatic anatomy and function of the mouse tail, 5 mg/kg hucMSC-Ex was injected subcutaneously into the area of edema. In vivo fluorescent imaging results showed that Dir labeled hucMSC-Ex targeted injured tail 24 h post-injection (Figure 1D). Intradermal injections of methylene blue on day 42 after the primary operation resulted in drainage of the dye across the site of incision in the mice treated with hucMSC-Ex but not in those treated with PBS (Figure 1E). A marked reduction of the edema was observed in the hucMSC-Ex treated mice, which did not occur in the PBS treated mice (Figure 1E). Tail volume was increased significantly over time at 0, 1, and 2 cm distal to the incision site in the PBS treated mice with lymphedema (Figure 1F). On the contrary, tail volume was decreased in the hucMSC-Ex treated mice at 0 cm distal to the incision site on week 5, 1 cm distal to the incision site on week 4, and 2 cm distal to the incision site on week 4 (Figure 1F). Then the mouse tails were harvested and analyzed for lymphatic architecture and formation of neolymphangiogenesis. In histological sections, hucMSC-Ex therapy led to a relatively thinner dermis and epidermis while better preserving the epidermal/dermal junction (Figure 2A). Regenerated lymphatic vessel endothelial hyaluronan receptor 1 (LYVE-1) positive lymphatic vessels, which are specific makers of lymphangiogenesis, were found at the site of surgical preparation in hucMSC-Ex group. The number of lymphatics and the lymphatic area increased in the hucMSC-Ex treated mice (Figure 2B). These findings suggest that hucMSC-Ex can promote lymphangiogenesis and reduce lymphedema in vivo.

**Figure 1.**
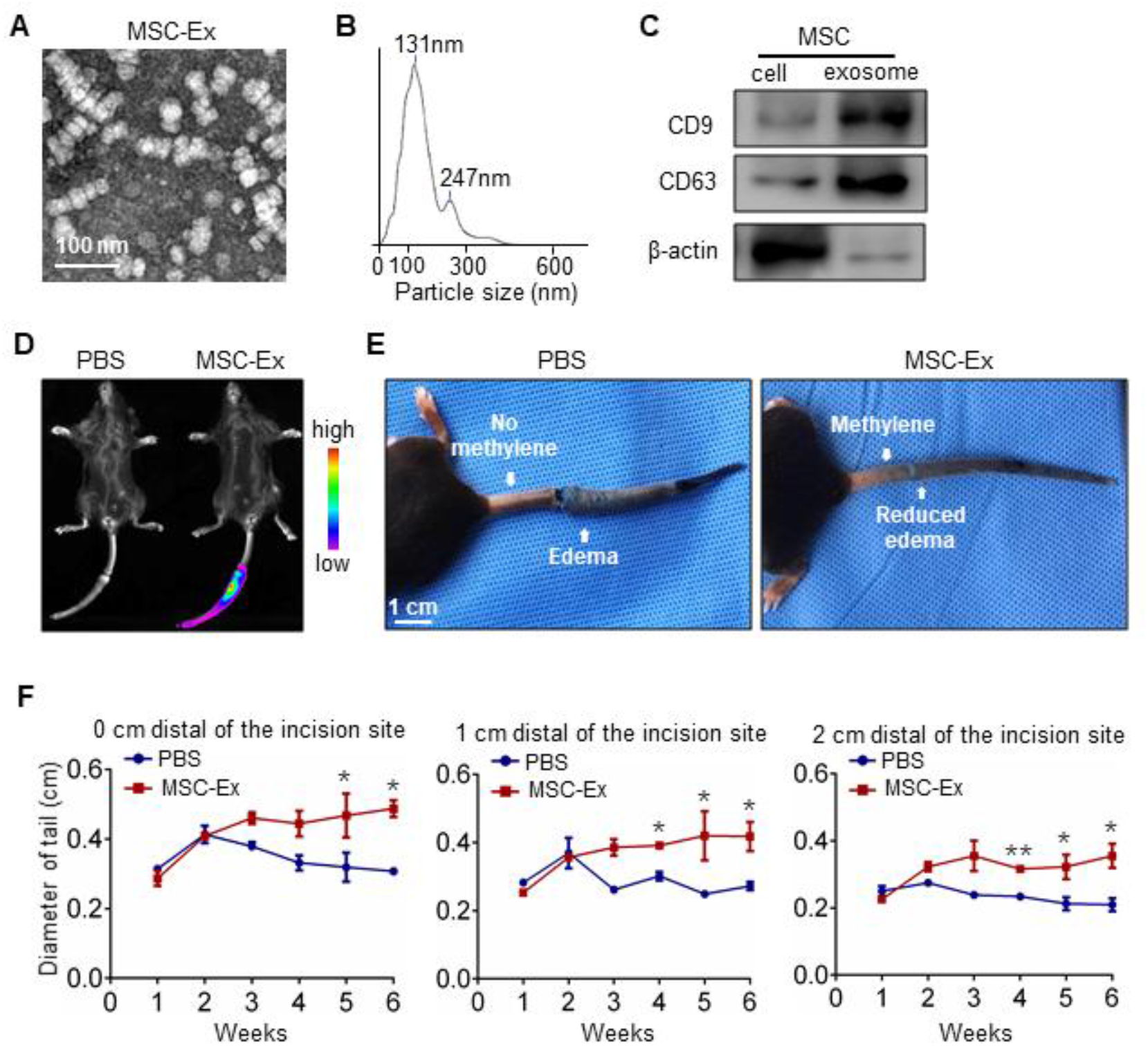
hucMSC-Ex treatment restores lymphatic function and reduces lymphedema in vivo. A Transmission electron microscopy (TEM) analysis of hucMSC-Ex. Scale bar, 100nm. B Nanoparticle tracking analysis (NTA) of the hucMSC-Ex diameter. C Western blot analysis showing exosome markers CD9 and CD63 in hucMSC-Ex. D Distribution of Dir labeled hucMSC-Ex in mouse tail with lymphatic ablation microsurgery after subcutaneous injections according to in vivo fluorescent imaging 24 h post-treatment. E hucMSC-Ex restored the lymphatic drainage functionality in a mice lymphedema model. Representative images showing intradermal injection of methylene blue on day 42 after the primary operation resulted in transportation of the dye across the incision site in mice receiving local application of hucMSC-Ex. Mice receiving no hucMSC-Ex showed no visible transportation of the dye across the incision site (n = 6). F Quantitation of mouse tail lymphedema volume after lymphatic ablation microsurgery with PBS or hucMSC-Ex treatment. Assessment of the circumference directly at the incision site at 0 cm, 1 cm, and 2 cm distal of the incision site showed progressively reduced edema formation in mice treated with hucMSC-Ex (n = 6, * P < 0.05, ** P < 0.01); data are presented as means ± SEM, Mann-Whitney test.

**Figure 2.**
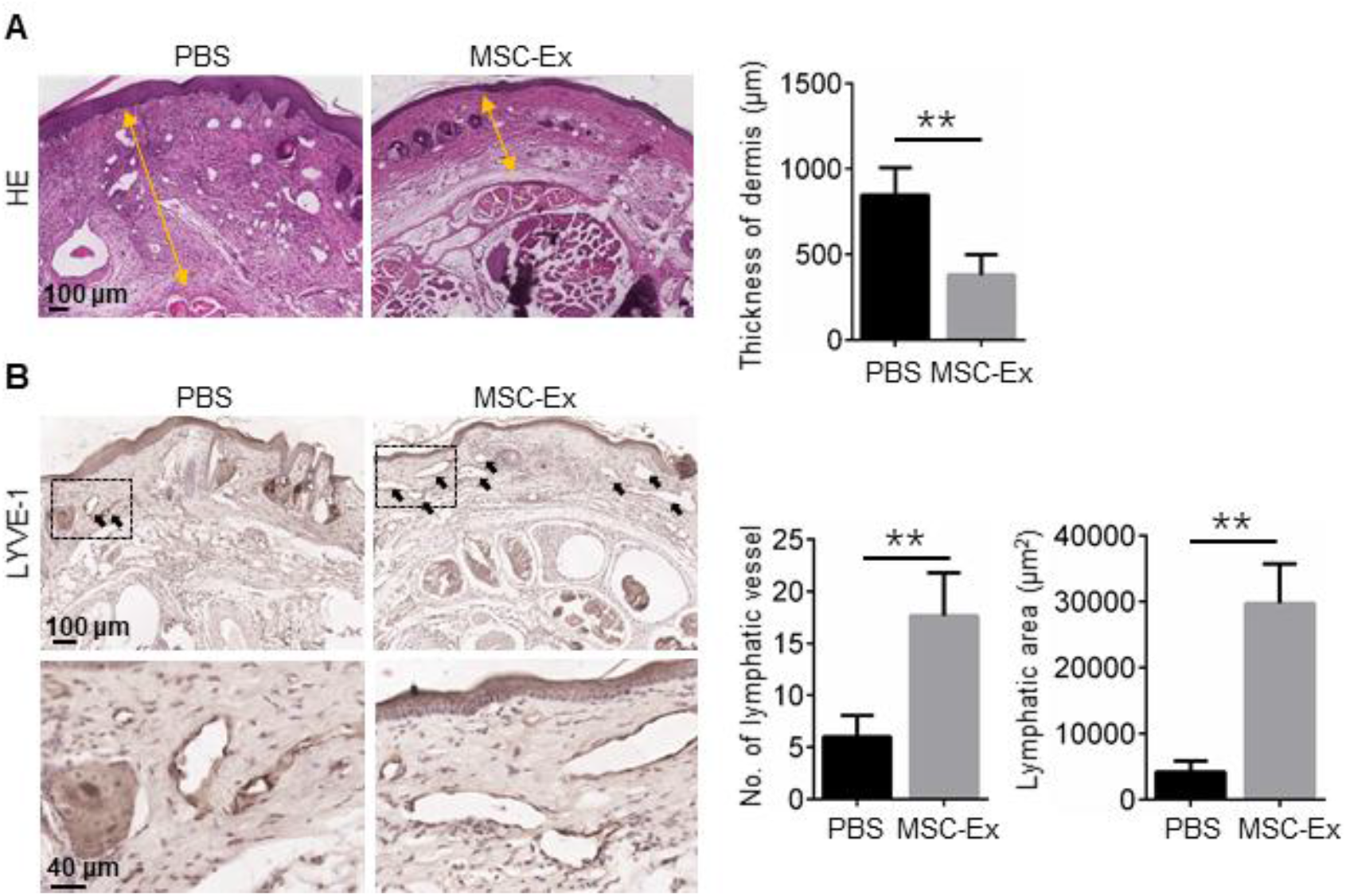
hucMSC-Ex treatment improves tail anatomy and regenerates lymphatic vessels in vivo. A Representative histology of mouse tail harvested on day 42 comparing samples from mice treated with PBS or hucMSC-Ex after lymphatic ablation microsurgery. Cutaneous dimension is indicated by yellow arrows. Dermal thickness was quantified on day 42 in tail skin (n = 6, ** P < 0.01). B Immunohistochemical staining for LYVE-1 positive regenerated lymphatic vessels. Number and area of LYVE-1 positive lymphatic vessels were quantified in the day 42 tail skin (n = 6, ** P < 0.01). For (A) and (B), data are presented as mean ± SEM; Mann-Whitney test.

### hucMSC-Ex Enhances Lymphangiogenesis In Vitro

To examine the uptake of hucMSC-Ex, the lipophilic dye PKH26 was used to label hucMSC-Ex, and this compound was incubated with HDLECs for different times. Fluorescent microscopic imaging showed that hucMSC-Ex was found in HDLECs after 12 h of incubation, and that the quantity increased over time (Figure 3A). We then analyzed the effect of hucMSC-Ex treatment on the expression of the lymphatic marker LYVE-1 in HDLECs. Double immunofluorescence staining and western blot analysis indicated that CD63 positive hucMSC-Ex treatment promoted LYVE-1 expression time-dependently in HDLECs (Figure 3B and 3C). We then investigated the prolymphangiogenic ability of hucMSC-Ex by regulating HDLEC migration, tube formation, and proliferation. hucMSC-Ex dose-dependently promoted HDLEC migration (Figure 3D and 3E). The Matrigel tube formation assay also documented hucMSC-Ex induced tube formation, whereas PBS did not affect tube formation (Figure 3D and 3F). Furthermore, the CCK-8 assay showed that HDLEC proliferation was promoted by hucMSC-Ex, whereas comparable doses of PBS were inactive (Figure 3G). The results demonstrate that hucMSC-Ex can enhance lymphangiogenesis by promoting HDLECs proliferation, migration, and tube formation.

**Figure 3.**
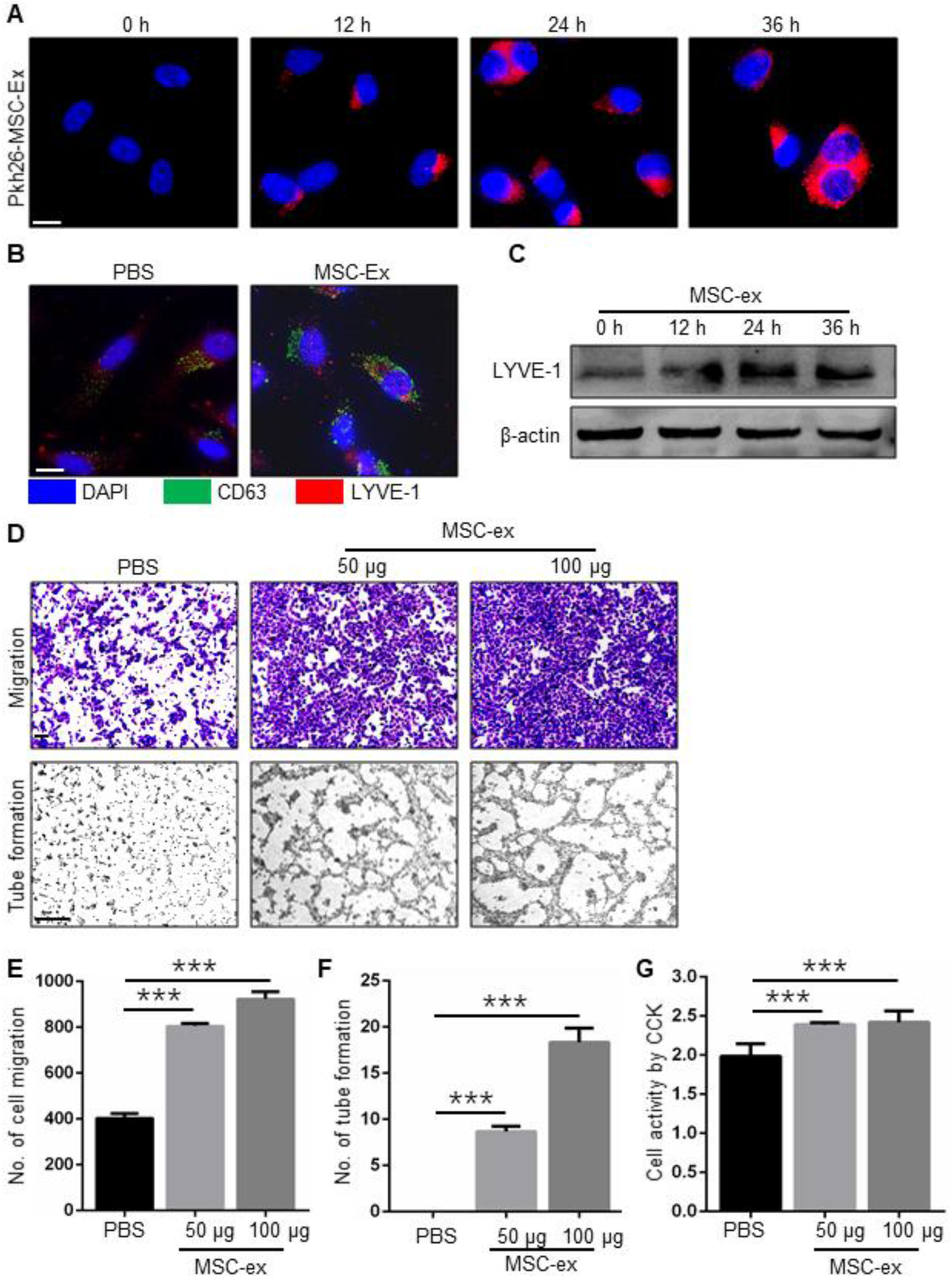
hucMSC-Ex exhibits dose-dependent effect on lymphangiogenesis in vitro. A Location of PKH26-labeled hucMSC-Ex (red) in HDLECs detected by imaging flow cytometry at 12, 24, and 36 h post hucMSC-Ex incubation. Scale bars, 10 μm (n = 3). B Representative immunofluorescent images of CD63 and LYVE-1 expression in PBS or hucMSC-Ex treated HDLECs at 24 h (n = 3). Original magnification 400×, Scale bars, 10 μm. C LYVE-1 expression quantified by western blot. hucMSC-Ex dose-dependently improved LYVE-1 expression in HDLECs. D Representative images of HDLEC migration and tube formation assays. HDLECs were treated with different concentrations of hucMSC-Ex. 50 and 100 μg hucMSC-Ex have prolymphangiogenic activity. Original magnification for HDLEC migration 100×; Original magnification for HDLEC tube formation 40×. Scale bars, 100 μm. E-G Quantitative analysis of HDLECs subjected to hucMSC-Ex treatment in migration, tube formation, and proliferation assays. For (E) (n = 3, *** P < 0.001). For (F) (n = 3, *** P < 0.001), and (G) (n = 5, *** P < 0.001), data are presented as means ± SEM; comparisons with the control groups were made using one-way ANOVA test followed by Dunn’s multiple comparisons test.

### hucMSC-Ex Improves Ang-2 mRNA and Protein Level in HDLECs

Lymphangiogenesis is controlled by a number of growth factors. (Alitalo, 2011, Yoshimatsu, Miyazaki et al., 2016) To investigate the components that have a role in controlling hucMSC-Ex effects on therapeutic lymphangiogenesis, we screened lymphangiogenic factors in hucMSCs-Ex and hucMSCs. Results of the western blot analyses showed that Ang-2 (but not VEGFR3 or Prox1) was expressed in hucMSCs and hucMSC-Ex (Figure 4A; Figure EV1). Treatment with hucMSC-Ex induced Ang-2 expression in HDLECs with about a 60% increase in the mRNA level at 48 h (Figure 4B) by quantitative reverse transcription polymerase chain reaction (qRT-PCR) testing, and about a 3-fold increase in protein levels at 24 h (Figure 4C) compared with PBS controls, determined by western blot analyses. Immunofluorescence staining also confirmed that CD63 positive hucMSC-Ex treatment significantly increased Ang-2 protein in HDLECs at 24 h (Figure 4D). Thus, hucMSC-Ex may first increase Ang-2 protein by exosomal transfer and then induce Ang-2 mRNA expression.

**Figure4.**
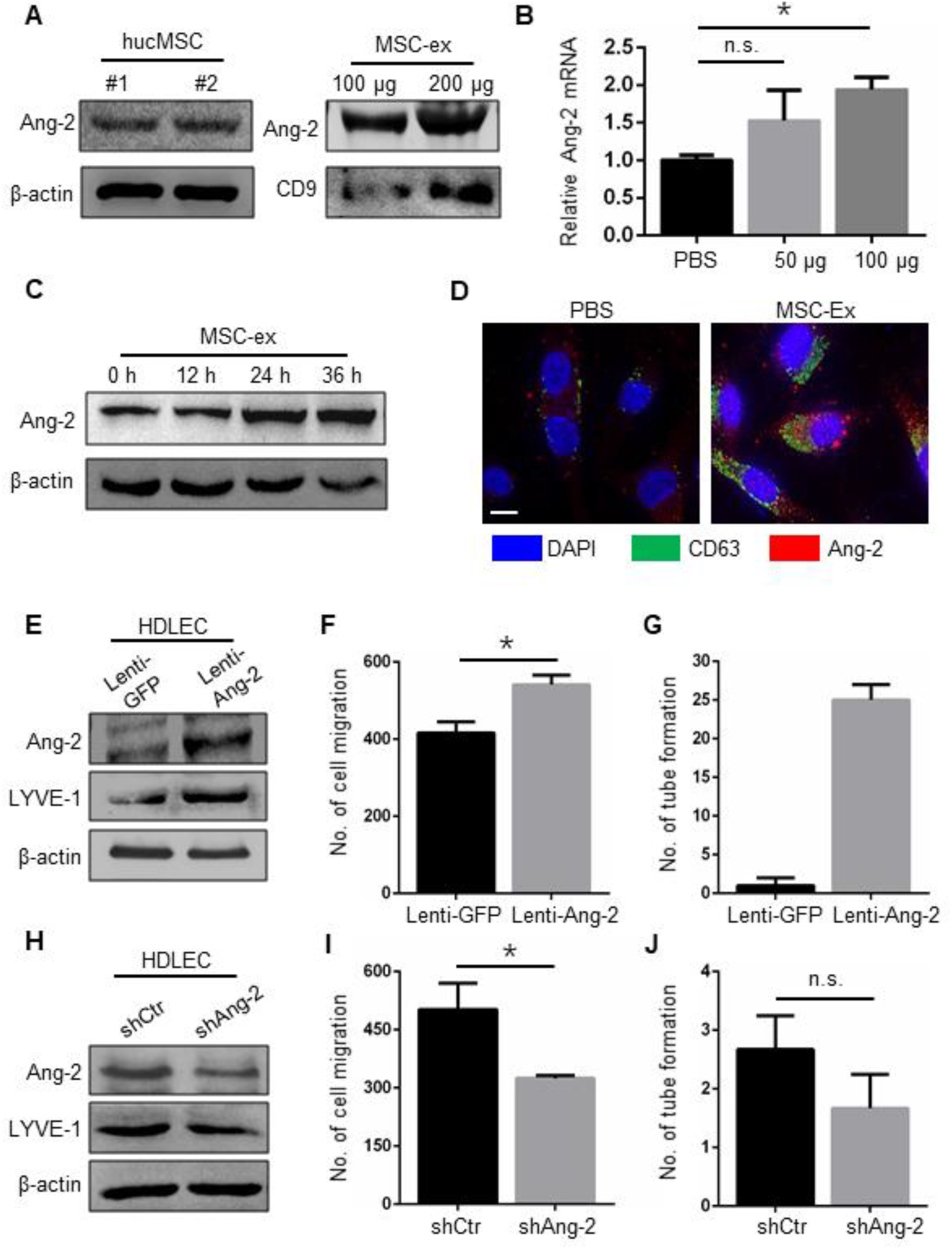
hucMSC-Ex derived Ang-2 is transferred to HDLECs in vitro. A Western blot analysis of Ang-2 expression in hucMSCs and hucMSC-Ex. Ang-2 protein was highly expressed in hucMSCs and hucMSC-Ex. B qRT-PCR analysis measured the relative transcripts of Ang-2 in HDLECs after the treatment of hucMSC-Ex; (n = 6, n.s. = not significant; * P < 0.05). C hucMSC-Ex treatment time-dependently increased Ang-2 protein in HDLECs. D Representative immunofluorescent images of CD63 and Ang-2 expression in PBS or hucMSC-Ex treated HDLECs at 24 h (n = 3). Scale bars, 10 μm. E Western blot analysis of Ang-2 and LYVE-1 expression in Lenti-GFP and Lenti-Ang-2 transfected HDLECs. Ang-2 and LYVE-1 expression were increased in Lenti-Ang-2 transfected HDLECs compared with which in Lenti-GFP. F-G Quantification of HDLEC migration, tube formation, and proliferation. Ang-2-overexpression promoted HDLEC migration (F) (n = 4, * P < 0.05) and tube formation (G) (n = 5, ** P < 0.01). H Western blot analysis of Ang-2 and LYVE-1 expression in control shRNA (shCtr), and shAng-2 transfected HDLECs. Ang-2 and LYVE-1 expression were decreased in shAng-2 transfected HDLECs compared with which in shCtr. I-J Quantification of HDLEC migration and tube formation. Ang-2 knockdown reduced HDLEC migration (I) (n = 4, * P < 0.05) and tube formation (J) (n = 5, n.s. = not significant). For (F) to (J), data are presented as means ± SEM; Mann-Whitney test.

Ang-2 play a crucial role in lymphangiogenesis by regulating lymphatic specification, interendothelial cell-cell junctions, and sprouting.(Alitalo, 2011, Yoshimatsu et al., 2016) To provide further evidence of hucMSC-Ex derived Ang-2 in lymphangiogenesis, we determined whether Ang-2 is involved in HDLECs lymphangiogenesis by regulating LYVE-1 expression, cell migration, tube formation, and proliferation. Overexpression of Ang-2 with Lenti-Ang-2 significantly upregulated LYVE-1 expression (Figure 4E), promoted HDLEC migration (Figure EV2A; Figure 4F) and tube formation (Figure EV2A; Figure 4G). Conversely, inhibition of Ang-2 expression by shRNA decreased LYVE-1 expression (Figure 4H) and HDLEC migration (Figure EV2B; Figure 4I), whereas tube formation was not reduced further because there was minimal tube formation in the parental cells (Figure EV2B; Figure 4J). Furthermore, HDLEC proliferation was promoted by Lenti-Ang-2 transfection compared to the Lenti-GFP controls and it was reduced by shAng-2 compared with ctr-shRNA (Figure EV3). These results indicate that Ang-2 can induce lymphangiogenesis by promoting HDLEC migration, tube formation, and proliferation in vitro.

### Ang-2 Is Critical in hucMSC-Ex Mediated Prolymphangiogenesis Effects In Vitro

To determine whether Ang-2 is essential for hucMSC-Ex to affect lymphangiogenesis, we compared the role of hucMSC-Ex with Ang-2 overexpression (Ex^Ang-2^) or knockdown (Ex^shAng-2^) in the regulation of LYVE-1 expression, HDLEC migration, tube formation, and proliferation. After Lenti-Ang2 transfection, Ang-2 was overexpressed in hucMSCs and hucMSC-Ex (Ex^Ang-2^) (Figure 5A). In comparison with Ex^GFP^, Ex^Ang-2^ significantly increased LYVE-1 levels (Figure 5B). Moreover, Ex^Ang-2^ promoted HDLEC migration, tube formation, and proliferation compared with Ex^GFP^ (Figure EV4A; Figure 5E). After knockdown with Ang-2 shRNA, Ang-2 was decreased in hucMSCs and hucMSC-Ex (Ex^shAng-2^) (Figure 5C). In contrast, improved Ang-2 levels by Ex^shCtr^, Ex^shAng-2^ reduced LYVE-1 expression in HDLEC (Figure 5D). Consistently, HDLEC migration, tube formation, and proliferation were impaired compared with that which occurred after Ex^shCtr^ (Figure EV4B; Figure 5F). These data demonstrate that Ang-2 has a critical role in hucMSC-Ex mediated lymphangiogenesis.

**Figure 5.**
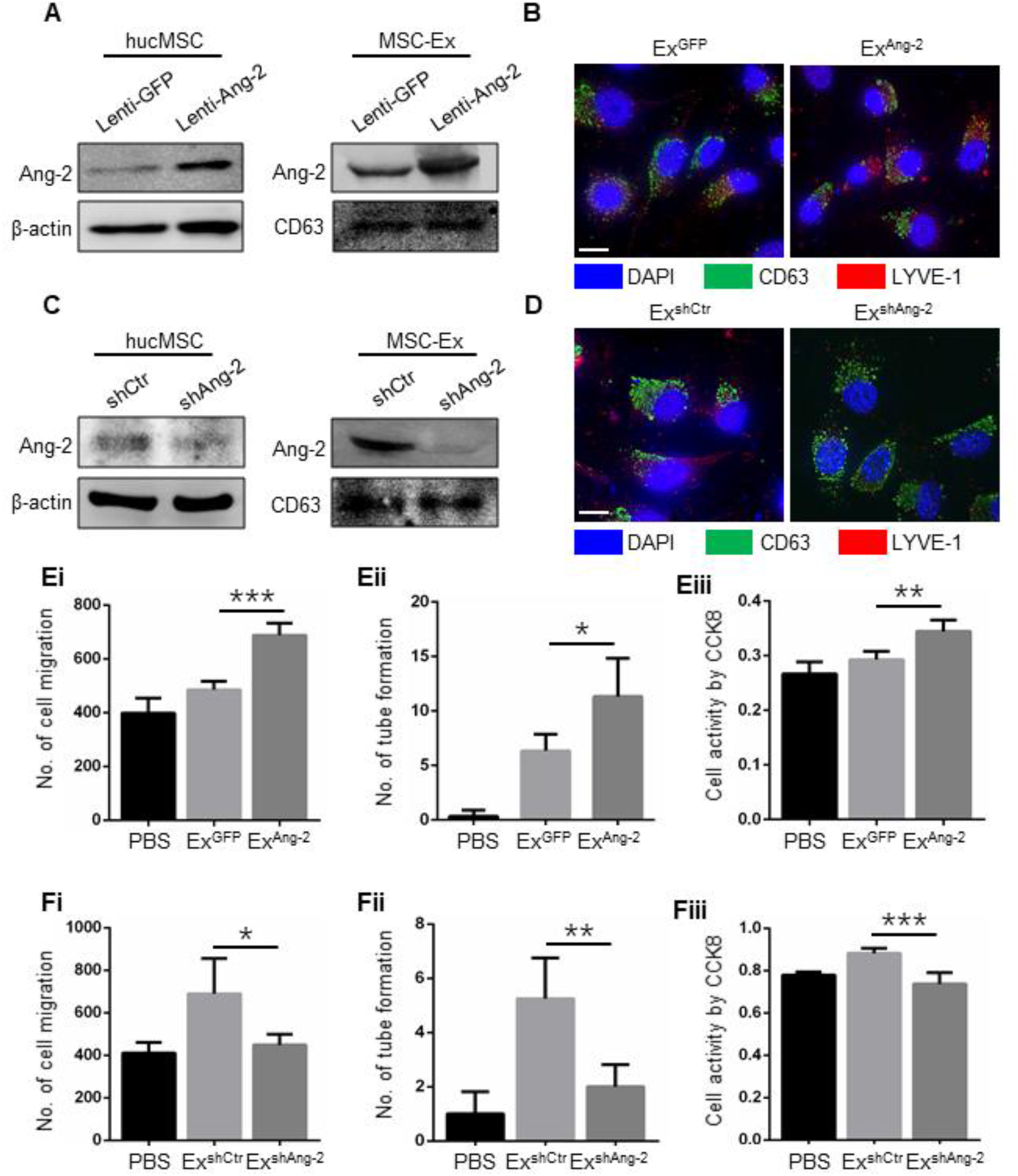
hucMSC-Ex derived Ang-2 promotes HDLEC lymphangiogenesis in vitro. A Western blot analysis of Ang-2 expression in Lenti-GFP and Lenti-Ang-2 transfected hucMSC and hucMSC-Ex. Ang-2 was overexpressed in Lenti-Ang-2 modified hucMSC-Ex (Ex^Ang-2^) compared with Ex^GFP^. B Representative immunofluorescent images of CD63 and LYVE-1 expression in Ex^GFP^ or Ex^Ang-2^ treated HDLECs at 24 h. Ex^Ang-2^ significantly increased LYVE-1 expression in HDLECs compared with Ex^GFP^ (n = 3). Scale bars, 10 μm. C Western blot analysis of Ang-2 expression in shCtr and shAng-2 transfected hucMSC and hucMSC-Ex. Ang-2 was knockdown in shAng-2 modified hucMSC-Ex (Ex^shAng-2^) compared with Ex^shCtr^. D Representative immunofluorescent images of CD63 and LYVE-1 expression in Ex^GFP^ or Ex^shAng-2^ treated HDLECs at 24 h. Ex^shAng-2^ significantly decreased LYVE-1 expression in HDLECs compared with Ex^shCtr^ (n = 3). Scale bars, 10 μm. E-F Quantification of HDLEC migration, tube formation, and proliferaion. Ex^Ang-2^ further improved HDLEC migration (Ei) (n = 5, *** P < 0.001), tube formation (Eii) (n = 3, * P < 0.05), and proliferation (Eiii) (n = 4, ** P < 0.01) in comparison with Ex^GFP^, which has a finite effect. In contrast, HDLEC migration (Fi) (n = 4, * P < 0.05), tube formation (Fii) (n = 4, ** P < 0.01), and proliferation (Fiii) (n = 4, *** P < 0.001) promotion by Ex^shCtr^ was diminished by Ex^shAng-2^. For (E) to (F), data are presented as means ± SEM; comparisons with the control groups were made using the one-way ANOVA test followed by Dunn’s multiple comparisons test.

### Ang-2 Enhanced Lymphangiogenesis via Activation of Prox1, VEGFR3/phospho-Akt Expression

The exact role of Ang-2 in lymphangiogenesis is still unclear. To corroborate hucMSC-Ex effects on lymphangiogenesis, we examined the regulation of hucMSC-Ex derived Ang-2 on Prox1, VEGFR3, and phospho/total-Akt (p-Akt/t-Akt) protein in HDLECs. First, hucMSC-Ex treatment increased HDLEC expression of Prox1, VEGFR3, and p-Akt, respectively, in a time-dependent fashion (Figure 6A). Next, we found that expression of Prox1, VEGFR3, and p-Akt in HDLECs was increased by Lenti-Ang-2 compared with Lenti-GFP and decreased by shAng-2 compared with shCtr (Figure 6B). Then, we confirmed that Ex^Ang-2^ improved the promotion of Ex^GFP^ on Prox1, VEGFR3, and p-Akt protein in HDLECs (Figure 6C and 6D), while Ex^shAng-2^ diminished the promotion of Ex^shCtr^ on Prox1, VEGFR3, and p-Akt protein in those cells (Figure 6E and 6F). These results indicated that Ang-2 can regulate Prox1, VEGFR3, and p-Akt expression in HDLECs, that have been reported to play an important role in LECs differentiation and lymphangiogenesis. Thus hucMSC-Ex derived Ang-2 may promote HDLEC proliferation, migration, and tube formation via regulating expression of Prox1 and VEGFR3/p-Akt.

**Figure 6.**
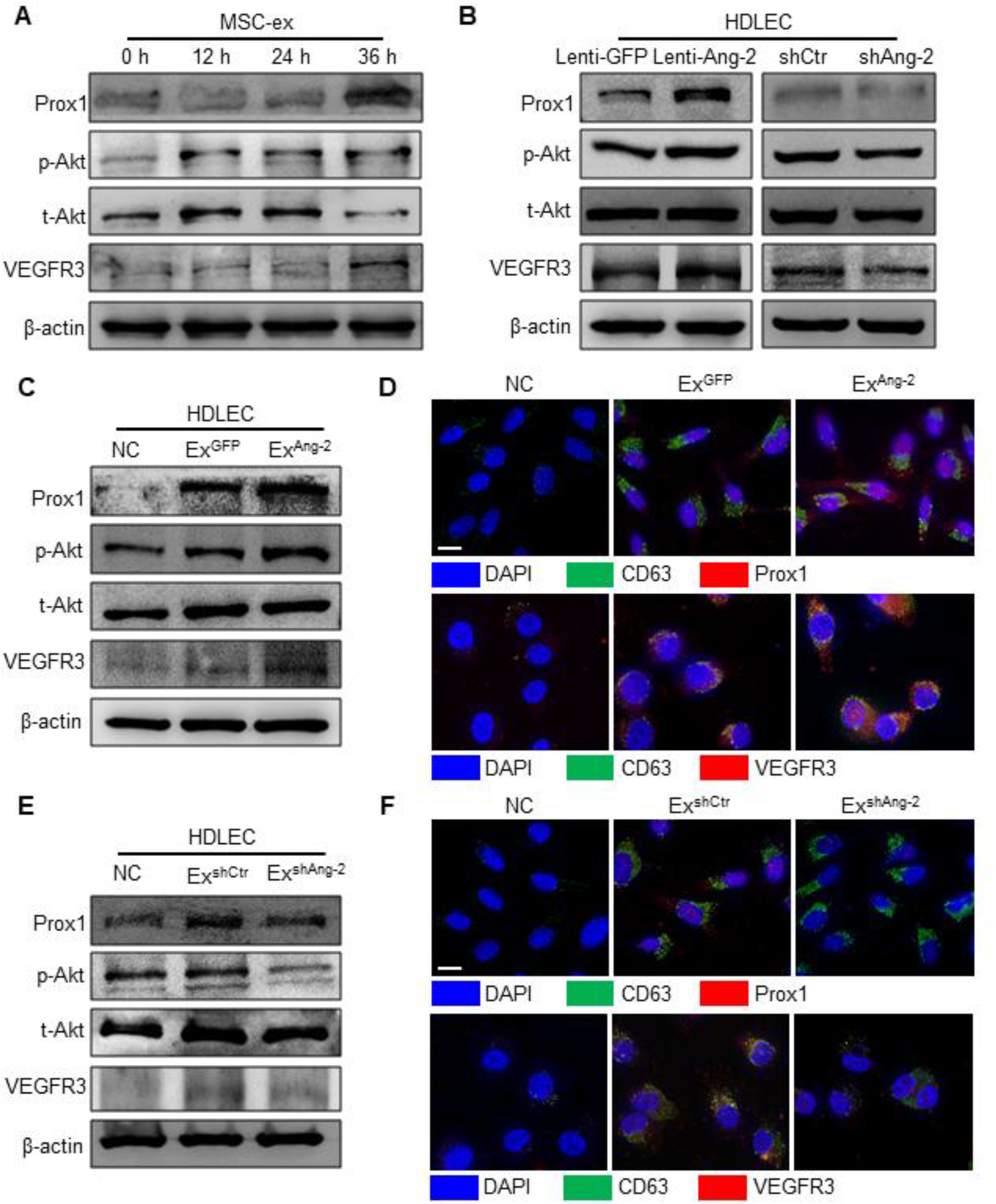
hucMSC-Ex derived Ang-2 induces Prox1 and VEGFR3/phospho-Akt expression in HDLECs. A Western blot analysis of Prox1, VEGFR3, phospho-Akt (p-Akt) and total-Akt (t-Akt) protein in HDLECs. Expression of Prox1, VEGFR3, and Akt phosphorylation were time-dependently increased in hucMSC-Ex treated HDLECs. B Western blot analysis of Prox1, VEGFR3, p-Akt and t-Akt protein in Lenti-GFP-, Lenti-Ang-2-, shCtr-, and shAng-2 transfected HDLECs. Expression of Prox1, VEGFR3, and p-Akt in HDLECs was increased by Lenti-Ang-2 compared with Lenti-GFP and decreased by shAng-2 compared with shCtr. C Western blot analysis of Prox1, VEGFR3, p-Akt and t-Akt protein in Ex^GFP^ and Ex^Ang-2^ treated HDLECs. Ex^Ang-2^ improved the promotion of Ex^GFP^ on Prox1, VEGFR3, and p-Akt in HDLECs. D Representative immunofluorescent images of CD63/Prox1 and CD63/VEGFR3 expression in Ex^GFP^ and Ex^Ang-2^ treated HDLECs at 24 h. Expression of Prox1 and VEGFR3 in HDLECs was increased by Ex^Ang-2^ compared with Ex^GFP^ (n = 3). Scale bars, 10 μm. E Western blot analysis of Prox1, VEGFR3, p-Akt and t-Akt protein in Ex^shCtr^ and Ex^shAng-2^ treated HDLECs. Ex^shAng-2^ diminished the promotion of Ex^shCtr^ on Prox1, VEGFR3, and p-Akt protein in HDLECs. F Representative immunofluorescent images of CD63/Prox1 and CD63/VEGFR3 expression in Ex^shCtr^ and Ex^shAng-2^ treated HDLECs at 24 h. Expression of Prox1 and VEGFR3 in HDLECs was decreased in Ex^shAng-2^ compared with which in Ex^shCtr^ (n = 3). Scale bars, 10 μm.

### hucMSC-Ex Derived Tie2 Is Transferred to HDLECs In Vitro

Ang-2 is a secreted ligand for the Tie2 receptor tyrosine kinase and Ang-2/Tie2 has been demonstrated to regulate lymphangiogenesis.(Yan, Jiang et al., 2012, Yoshimatsu et al., 2016) To further delineate the mechanism of hucMSC-Ex in lymphangiogenesis, we analyzed the expression of Ang-2 and Tie2 in hucMSC-Ex treated HDLECs. The results of these immunofluorescence studies showed that hucMSC-Ex increased Ang-2/Tie2 co-localization in HDLECs compared with PBS (Figure 7A). Additionally, hucMSC-Ex increased Tie2 expression in a time-dependent manner (Figure 7B). Interestingly, abundant Tie2 protein was found in hucMSC-Ex (Figure 7C). Neither knockdown nor overexpression of Ang-2 with Lenti-Ang-2 or shAng-2 had an effect on Tie2 expression (Figure EV5; Figure 7D). These results indicate that hucMSC-Ex promote lymphangiogenesis by delivering exosomal Ang-2/Tie2 protein to HDLECs inducing Prox1 and VEGFR3/p-Akt proteins expression (Figure 7E).

**Figure 7.**
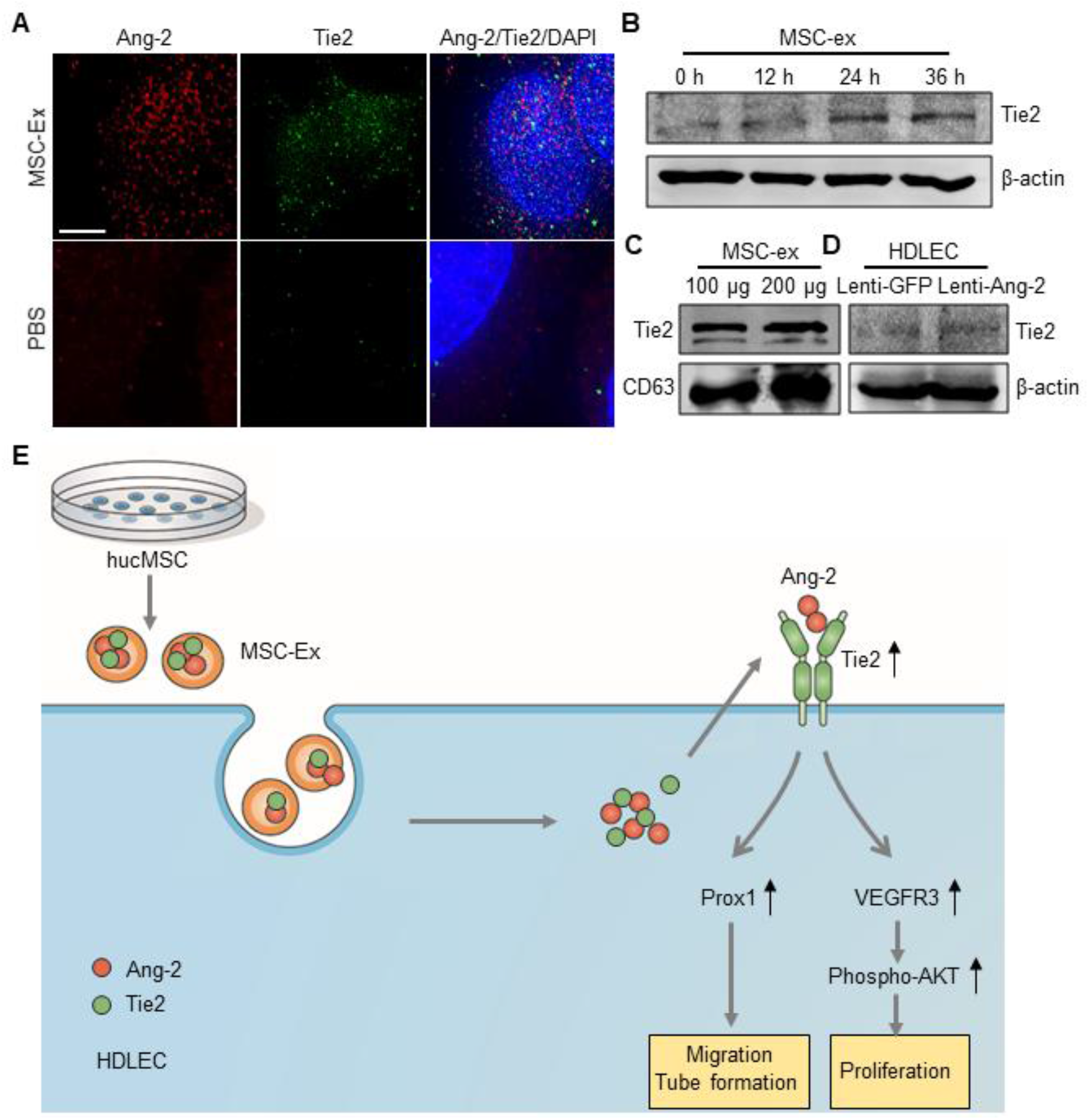
hucMSC-Ex derived Tie2 is transferred to HDLECs in vitro. A Representative immunofluorescent images of Ang-2 and Tie2 expression in PBS or hucMSC-Ex treated HDLECs at 24 h. hucMSC-Ex increased Ang-2 and Tie2 expression in HDLECs compared with PBS (n = 3). Scale bars, 5 μm. B hucMSC-Ex time-dependently increased Tie2 protein in HDLECs. C Western blot analysis of Tie2 protein in hucMSC-Ex. D Western blot analysis of Tie2 expression in Lenti-GFP and Lenti-Ang-2 transfected HDLECs. Lenti-Ang-2 transfection has no effect on Tie2 expression in HDLECs compared with Lenti-GFP control. E Summary of the new knowledge based on our hypothesis. Exosomes from hucMSCs induced HDLEC migration, tube formation, and proliferation by delivering exosomal Ang-2/Tie2 protein to HDLECs inducing Prox1 and VEGFR3/p-Akt signaling.

## Discussion

The role and mechanism of hucMSC-Ex in the process of recovery from lymphedema is largely unknown. In the present study we demonstrated for the first time that hucMSC-Ex can promote lymphangiogenesis in vitro and in vivo. The administration of hucMSC-Ex in vivo contributed to the regeneration of LYVE-1 positive lymphatic vessels and reduction of lymphedema, a prevalent and clinically relevant problem. Furthermore, exosomal Ang-2 plays an important role in hucMSC-Ex effects on lymphangiogenesis. Finally, we mechanistically supported exosomal Ang-2/Tie2 increases of HDLEC migration and tube formation via the Prox1 and VEGFR3/p-Akt signaling cascade. Based on these findings, we derived a schematic of our working hypothesis as to how hucMSC-Ex enhances lymphangiogenesis and reduces lymphedema. The findings extend our understanding of the function of hucMSC-Ex and suggest a new mechanism of Ang-2/Prox1 and Ang-2/VEGFR3 /p-Akt signaling in lymphangiogenesis.

The capacity of MSCs to secrete growth factors give them the potential to be an ideal therapy for lymphedema.(Schinkothe, Bloch et al., 2008, Takeda, Sowa et al., 2015) Clinical trials showed that MSC treatment was more effective for lymphedema than conservative compression therapy.(Hou et al., 2008, Maldonado et al., 2011) Exosomes are 30-150 nm diameter extracellular vesicles that deliver various cellular derived factors to target cells. Our previous studies reported that MSC derived exosomes can be engulfed by tissues locally or carried in the bloodstream to distant sites in the body. Here we found that Ang-2 and Tie2, two important lymphangiogenic factors, can be secreted by human MSCs without an external stimulus and were enriched in hucMSC-Ex. Other lymphangiogenic factors, such as Prox1 and VEGFR3, were not found. We thus investigated the role and mechanism of exosomal Ang-2 in lymphangiogenesis and lymphedema.

Lymphangiogenesis consists of a complex process that involves LEC proliferation, migration, and tube formation.(Conrad et al., 2009, Kajiya, Hirakawa et al., 2005, Robering, Weigand et al., 2018) Ang-2 controls lymphangiogenesis by adjusting lymphatic differentiation and sprouting throughout embryonic and neonatal development.(Dellinger, Hunter et al., 2008, Gale, Thurston et al., 2002) Deletion of Ang-2 resulted in defective remodeling of both collecting lymphatic vessels and capillaries, thereby resulting in lymphedema.(Gale et al., 2002) However, the exact role of Ang-2 in lymphangiogenesis is still unclear. We previously found that Ang-2 treatment cannot affect the proliferation of LECs.(Yan et al., 2012) In the present study, our data further suggested that HDLEC proliferation; migration and tube formation was promoted by Ang-2 overexpression with Lenti-Ang-2 transfection and was inhibited by Ang-2 knockdown. In addition, we confirmed that Ang-2 regulates the effect of hucMSC-Ex on HDLEC lymphangiogenesis, including proliferation, migration, and tube formation. Our findings establish that Ang-2 plays a critical role in hucMSC-Ex mediated lymphangiogenesis.

In this study, we found CD63-positive hucMSC-Ex located in the cytoplasm of HDLECs in vitro. Exosomal transfer of Ang-2 protein increased the intracellular Ang-2 of HDLECs. Furthermore, hucMSC-Ex also induced endogenous Ang-2 expression. Therefore, hucMSC-Ex uptake not only increased Ang-2 protein but also promoted Ang-2 mRNA in HDLECs. In addition, hucMSC-Ex induced about a 3-fold increase in protein levels at 24 h and about a 60% increase in mRNA levels at 48 h. Thus, Ang-2 induction in HDLECs mainly depends on the action of exosomal Ang-2 rather than endogenous Ang-2.

Among the factors that drive the developmental differentiation of lymphatic structures from venous endothelium, Prox1 is the master transcriptional regulator for the early steps of LECs differentiation from the embryonic veins, and it remains required for lymphatic identity.(Yang & Oliver, 2014, Yoshimatsu et al., 2016) VEGFR3 is the major tyrosine kinase receptor that leads to phosphorylation of the serine kinases Akt, which in turn promote LEC proliferation, migration, and survival.(Yoshimatsu et al., 2016, Zheng, Aspelund et al., 2014) To address the mechanism of Ang-2 signaling, we examined the expression of lymphangiogenic factors such as Prox1, VEGFR3, and p-Akt/t-Akt in hucMSC-Ex treated- and Ang-2 modified HDLECs. Our results showed that hucMSC-Ex derived Ang-2 induced Prox1, VEGFR3, and p-Akt expression. Ang-2 regulates lymphatic development through interaction with receptor tyrosine kinase Tie2.(Yoshimatsu et al., 2016) Here we showed that exosomal Tie2 was transferred to HDLECs. Thus hucMSC-Ex may enhance lymphangiogenesis via exosomal transfer of Ang-2 and Tie2. Based on these findings, we provide a schematic of our hypothesis as to how hucMSC-Ex increases lymphatic regeneration during lymphedema.

In conclusion, our data demonstrate that hucMSC-Ex has a previously unappreciated prolymphangiogenic role in lymphedema therapy. Our work also provides compelling evidence that exosomal Ang-2 is essential for the lymphatic regeneration during lymphedema. Exosomal Ang-2 and Tie2 promote lymphangiogenesis through upregulating Prox1 and VEGFR3/p-Akt expression in HDLECs, which may provide a new therapeutic modality for boosting the efficacy of therapeutic lymphangiogenesis.

## Materials and Methods

### Animals

The animal experiments were approved by the ethics committee of Jiangsu University, and the methods were carried out in accordance with the approved guidelines.

### Cell Culture

hucMSCs were isolated and characterized as described previously and maintained in α-MEM (Gibco, New York, USA) containing 10% fetal bovine serum (FBS) (ExCell Bio, Taicang, China) at 37°C with 5% CO_2_. Human dermal lymphatic endothelial cells (HDLECs) were purchased from the Chinese Academy of Sciences and maintained in H-DMEM (Gibco) containing 10% FBS (ExCell Bio) at 37°C with 5% CO_2_. All cells were tested for mycoplasma contamination.

### Isolation and Characterization of hucMSC-Ex

The hucMSC-Ex were extracted and purified from hucMSCs as previously described.(Jiang et al., 2018) Briefly, hucMSCs were cultured in serum-free α-MEM (Gibco) for 48 h and condition medium was collected for hucMSC-Ex isolation. The protein concentration of the hucMSC-Ex was determined by using a BCA protein assay kit (CWBIO, Beijing, China). The final concentration of hucMSC-ex for treating lymphedema was 15 mg/kg. Exosomal markers CD9 and CD63 were determined by western blot. The morphology and size of the hucMSC-Ex were analyzed by transmission electron microscopy (FEI Tecnai 12, Philips, Netherlands) and nanosight tracking analysis (NanoSight, Amesbury, UK), respectively.

### Acquired Lymphedema by Lymphatic Ablation Microsurgery

Acquired lymphedema was surgically induced in C57BL/6J female mice (Laboratory animal center, Jiangsu University) tails using the thermal ablation technique for lymphatic trunks developed by Claudius Conrad.(Conrad et al., 2009) Briefly, the surgery was performed after general anesthesia with 8% (v/v%) chloral hydrate by intraperitoneal injection. A circumferential incision was made and a segmental band (2-mm wide) of skin 1 cm distal to the tail base was removed. To ensure that the ablation of the deeper lymphatics running alongside the major blood vessels had been successful, 5 μL of methylene blue was injected intradermally at the distal end of the tail, and any and all of the blue stained lymphatics at the site of the incision were excised under a surgical microscope. Care was taken to maintain the integrity of the major underlying blood vessels and tendons to avoid the tail distal to the incision did not become necrotic. After significant lymphatic edema developed within one week, the mice with lymphedema were randamly separated it into 2 groups: PBS (n=6), and MSC-Ex (n=6). 5 mg/kg of hucMSC-Ex was injected subcutaneously into the area of edema. PBS served as a control. Tail diameters at 0 (about 0.1 cm distal to the wound), 1, and 2 cm distal to the incision site were measured by observers blinded to the treatment using vernier caliper once a week after hucMSC-Ex or PBS administeration.

### hucMSCs Labeling and Tracking in HDLECs and Mouse Tail

For the in vivo tracking of hucMSC-Ex in mouse tail, 1,1-dioctadecyl-3,3,3,3-tetramethylindotricarbocyanine iodide (DiR) (Ruitai Biology, Shanghai, China) labeled hucMSC-Ex were injected subcutaneously in the area of edema and analyzed using a Maestro In Vivo Imaging System (CRI, MA, USA). In vivo spectral imaging from 690–850 nm was performed using an exposure time of 150 ms per image frame. To determine the in vitro distribution of hucMSC-Ex in HDLECs, PKH26 (Sigma, Merck, Germany) labeled hucMSC-Ex (PKH26-MSC-Ex) was incubated with the HDLECs at 37°C for different times and the fluorescent images were observed.

### Analysis of Lymphatic Drainage

To compare lymphatic vessel regeneration on day 42 between the mice treated with hucMSC-Ex and the controls, 5 μL of methylene blue was injected intradermally at the distal end of the tail and the lymphatic drainage of the dye was observed after 15 minutes. At the end of the experiment, the distribution of methylene blue staining in the mouse tails was observed and photographed using a camera (n=6) (Olympus, Japan). The field of view of the CCD detector were centered over the tail incision site.

### Histopathological Staining, Immunohistochemistry, and Immunofluorescence

At the end of the experiments, hucMSC-Ex or PBS treated mouse tails were surgically collected after general anesthesia with 8% (v/v%) chloral hydrate by intraperitoneal injection. The edematous tissues were gradually dehydrated, embedded in 10% paraffin, cut into 4-mm sections, and stained with H&E stain for light microscopy. The dermal lymphatic vessels in the mouse tails were identified using LYVE-1, an immunohistochemical marker of lymphatic endothelial cells. The tissue sections were then incubated with an anti-LYVE-1 monoclonal antibody (Abcam, Cambridge, UK) according to the manufacturer’s instructions. Signals were visualized using 3,3′-Diamino-benzidine tetrahydrochloride (Boster Biology, Wuhan, China). Immunofluorescence staining was performed using the following primary antibodies: anti-CD63 (Abcam), anti-LYVE-1 (Abcam), anti-Prox1 (Abcam), and anti-VEGFR3 (Bioworld, Nanjing, China). Negative controls with isotype IgG were run in parallel. Images were acquired using an inverted wide-filed fluorescence microscope (DeltaVision Elite, GE Healthcare Life Sciences).

### Cell Proliferation

HDLEC proliferation was assessed using a CCK-8 assay (Cell Counting Kit-8, Vazyme Biotech, Nanjing, China) as previously described. HDLECs were plated in 96-well plates in triplicate at 1×10^3^ cells per well and cultured in H-DMEM overnight at 37°C in 5% humidified CO_2_. After the medium was removed, HDLECs were washed with PBS and treated with hucMSC-Ex. Negative controls were prepared by PBS addition. The number of cells was measured by the absorbance (450nm) of reduced WST-8 (2-(2-methoxy-4-nitrophenyl)-3-(4-nitrophenyl)-5-(2,4-isulfophenyl)-2H-tetrazolium, monosodium salt) at the indicated time points.

### Cell Migration

HDLEC migration was examined using Transwell chambers with inserts of 8-µm pore size (Corning Costar, Corning, NY, USA) as described previously. The cells (5×10^4^ HDLECs) were suspended in serum-free media and plated onto the upper chamber for the migration assay following treatment with hucMSC-Ex or PBS, and media supplemented with 10% FBS was placed in the lower chamber. Migrated cells through the membrane to the lower surface were fixed, stained and counted after 8 h.

### Tube Formation

Matrigel tube formation was performed as previously described. Briefly, Matrigel (BD Biosciences, Bedford, MA, USA) was placed in 24-well plates (100 μL/well) and allowed to gel at 37°C for 30 minutes, before 6×10^4^ HDLECs that had been co-cultured with or without hucMSC-Ex for 36 h previously were seeded onto the coated wells and cultured in H-DMEM with 10% FBS at 37°C for 24 h under 5% CO_2_. and then HDLECs tube formation was observed.

### Quantitative RT-PCR

Total RNA was extracted from the HDLECs with the Trizol and a DNase (Invitrogen, Thermo fisher, MA, USA) treatment to remove any genomic DNA. cDNA was synthesized using oligo dT primers using SuperScript™ II RT kits according to the manufacturer’s instructions (Invitrogen). Quantitative PCR (qRT-PCR) was performed using FastStart SYBR Green (Roche) on a Lightcycler 480. mRNA expression relative to GAPDH mRNA expression was calculated using the delta-delta threshold cycle (^ΔΔ^CT) method. Primers to amplify Ang-2 were designed as following: ForTGGGATTTGGTAACCCTTCA, RevGGTTGGCTGATGCTGCTTAT.

### Western Blot

Cells were harvested and lysed in RIPA buffer. Equal amounts of cell lysates were loaded and separated on a SDS-PAGE gel. Transferred membranes were incubated with the primary antibodies against phospho-Akt (Cell Signaling Technology [CST], Danvers, MA, USA), Akt (Signalway Antibody LLC [SAB], College Park, MD, USA), Ang-2 (Abcam, Cambridge, UK), CD9 (Proteintech, USA), CD63 (Abcam, UK), LYVE-1 (Abcam, UK), Prox1 (Abcam, UK), Tie2 (Abcam, UK), and β-actin (Bioworld, Nanjing, China) overnight at 4°C. After incubation and washing with tris-buffered saline with 0.05% Tween-20 (TBST), membranes were challenged with horseradish peroxidase (HRP) conjugated goat anti-rabbit or goat anti-mouse antibodies, detected by the Amersham ECL detection system (GE Healthcare Life Sciences, Little Chalfont, UK)

### Lentiviral overexpression and knockdown of Ang-2

Lentiviruses expressing full-length human Ang-2 (Lenti-Ang-2), GFP alone (Lenti-GFP), Ang-2 shRNA (shAng-2), and control shRNA (shCtr) were constructed by Zhenjiang CarGene Biotechnology (Zhenjiang, Jiangsu, China). Full-length human Ang-2 cDNA (NCBI Reference Sequence: NM_001147) was inserted into pCDH-CMV-MCS-EF1-copGFP lentiviral vectors to generate Lenti-Ang-2 expression vectors. Ang-2 shRNA and control shRNA oligonucleotides were inserted into pPLK/GFP-Puro lentiviral vectors to generate Ang-2 shRNA expression vectors. Human Ang-2 shRNA oligonucleotide sequences are as follows: GGAAGAGCATGGACAGCATAG. The sequences of control shRNA are as follows: GTTCTCCGAACGTGTCACGTT. The lentivirus particals was produced by cotransfecting HEK293T cells with pPLK/GFP-Puro-Ang2 shRNA or pCDH-CMV-Ang2-EF1-copGFP, pMD2G and psPAX2 plasmids by using Lipofectamine 2000 (Invitrogen, Thermo fisher, MA, USA). After 72 h transfection, the virus containing supernatant was harvested and concentrated to a final vector titer of 8 × 10^7^ TU/ml. Lenti-Ang-2/Lenti-GFP and Lenti-Ang-2-shRNA/Lenti-control shRNA lentivirus were transduced into HDLECs and hucMSCs. The efficiency of Ang-2 overexpression and knockdown in HDLECs and hucMSCs was evaluated using Quantitative RT-PCR and western blot. For the preparation of Ang-2 modified hucMSC-Ex (Ex^Ang-2^ and Ex^shAng-2^), Lenti-Ang-2 and Ang-2-shRNA transfected hucMSCs were cultured in serum-free medium for 72 h. Exosomes were isolated for further research. For the preparation of Ang-2 modified HDLECs, Lenti-Ang-2 and Ang-2-shRNA transfected HDLECs were collected for further study.

### Statistics

The GraphPad Prism version 5.0 software package was used for statistical analysis. Differences between two groups at a single time point were compared using the Mann-Whitney test. For comparisons between multiple experimental groups, one-way ANOVA tests followed by Dunn’s multiple comparisons tests for post hoc analyses were used. All analyses were considered statistically significant at P < 0.05.

## Acknowledgments

This work was funded by the National Natural Science Foundation of China (81670549 and 81200312), the Natural Science Foundation of Jiangsu Province (BE2016717), Priority Academic Program Development of Jiangsu Higher Education Institutions (PAPD), and the Young Backbone Teacher Training Project of Jiangsu University.

## Author contributions

Y.Y., W.X., and H.Q. conceived experiments, analyzed results, and wrote the manuscript. T.Z., Z.Y., J.L., Y.C., Y.T., and H.S. performed experiments. Y.Y and H.Q. provided reagents, expertise, and feedback.

## The paper explained

### Problem

The therapeutic role and mechanisms of MSC-derived exosomes (MSC-Ex) in lymphedema is poorly understood.

### Results

human umbilical cord MSCs derived exosomes (hucMSC-Ex) treatment contributed to the regeneration of LYVE-1 positive lymphatic vessels and reduction of lymphedema in a mouse model of tail lymphedema. Following uptake, exosomal lymphangiogenic factors (angiopoietin (Ang)-2 and Tie2) are taken up by HDLECs and promoted HDLECs proliferation, migration, and tube formation in vitro. We also find that exosomal Ang-2 and Tie2 exert a prolymphangiogenic effect on HDLECs through upregulating Prox1 and VEGFR3/p-Akt expression.

### Impact

These results addressed a key undefined paracrine mechanism by which hucMSC-Ex promote lymphangiogenic effects, which have both scientific and clinical importance. It will lead to a deeper understanding of the mechanisms of hucMSC-Ex trafficking and function on one hand and therapeutic interventions on the other.

---

**Figure EV1.**
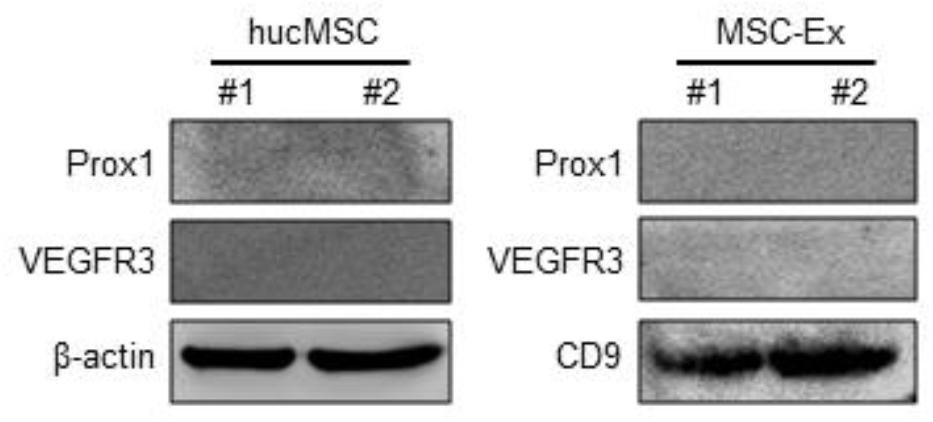
Western blot analysis of Prox1 and VEGFR3 protein in hucMSCs and hucMSC-Ex. Prox1 and VEGFR3 protein was not detected in hucMSCs and hucMSC-Ex.

**Figure EV2.**
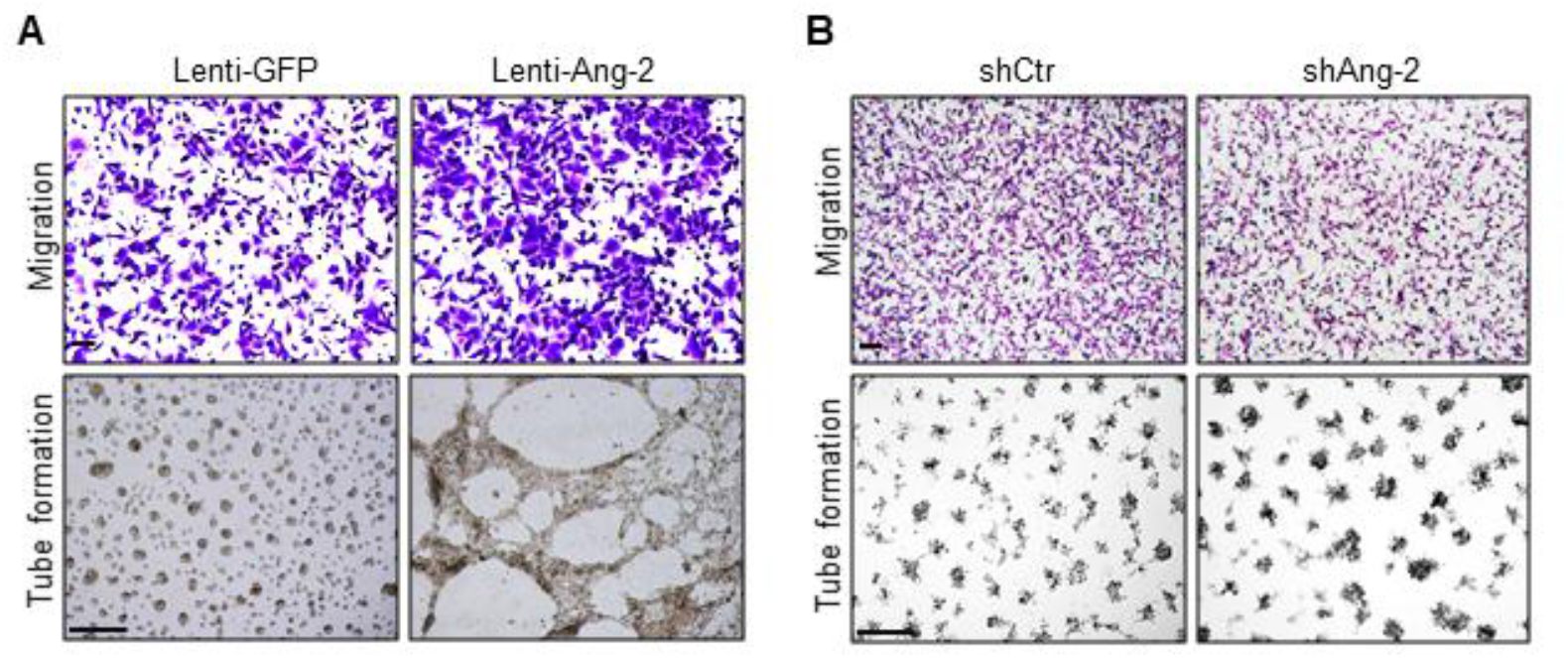
Representative images of cell migration and tube formation assays in Lenti-GFP, Lenti-Ang-2, shCtr, and shAng-2 transfected HDLECs. A Lenti-Ang-2 transfection increased HDLEC migration and tube formation compared with Lenti-GFP transfection. B shAng-2 transfection inhibited HDLEC migration and tube formation compared with shCtr. Original magnification for HDLEC migration 100×; Original magnification for HDLEC tube formation 40×. Scale bars, 100 μm.

**Figure EV3.**
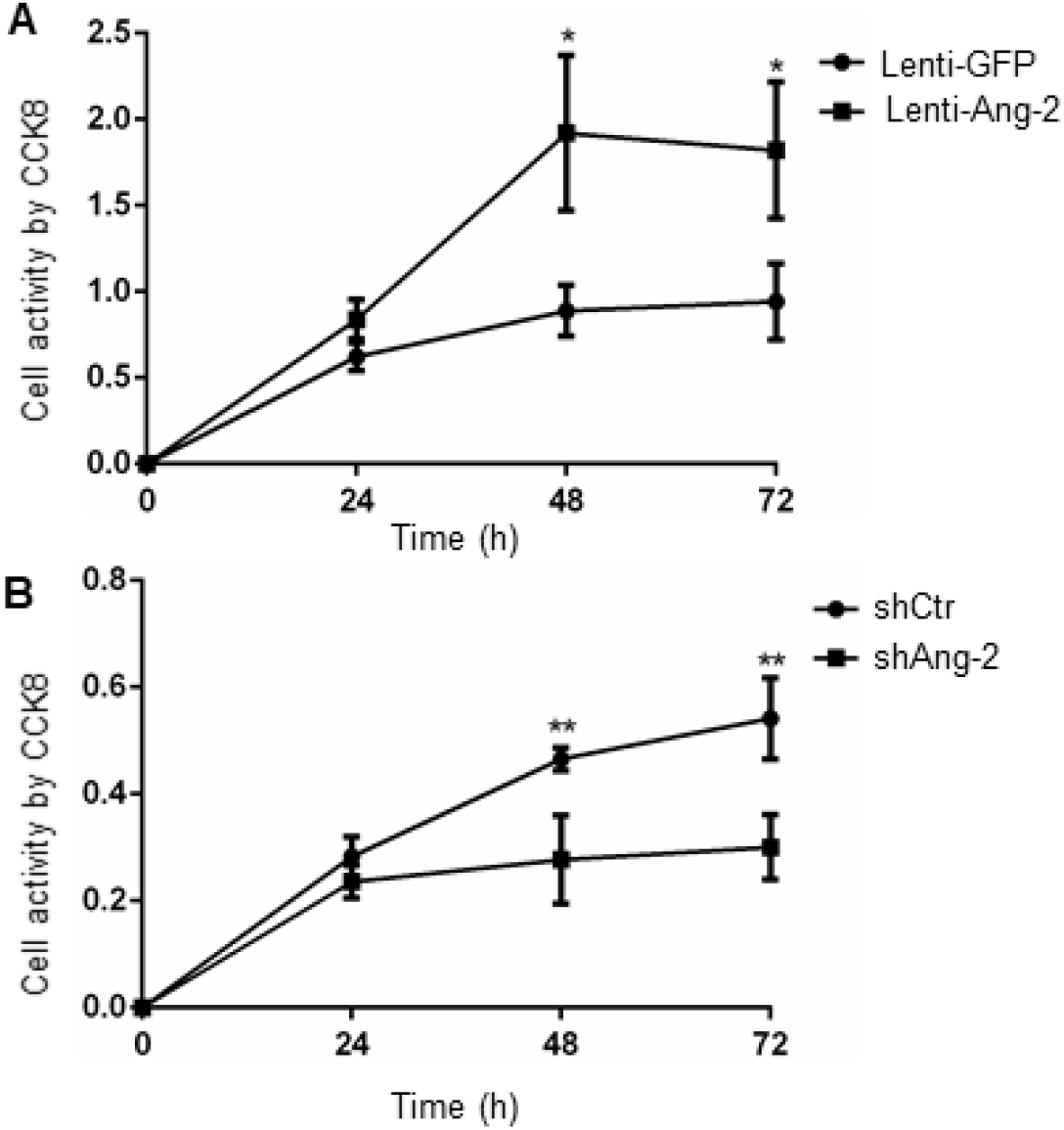
Quantification of HDLEC proliferation. A Ang-2- overexpression promoted HDLEC proliferation (n = 3, * p < 0.005). B Ang-2 knockdown reduced HDLEC proliferation (n = 4, ** p < 0.01). For (A) and (B), data are presented as means ± SEM; Mann-Whitney test.

**Figure EV4.**
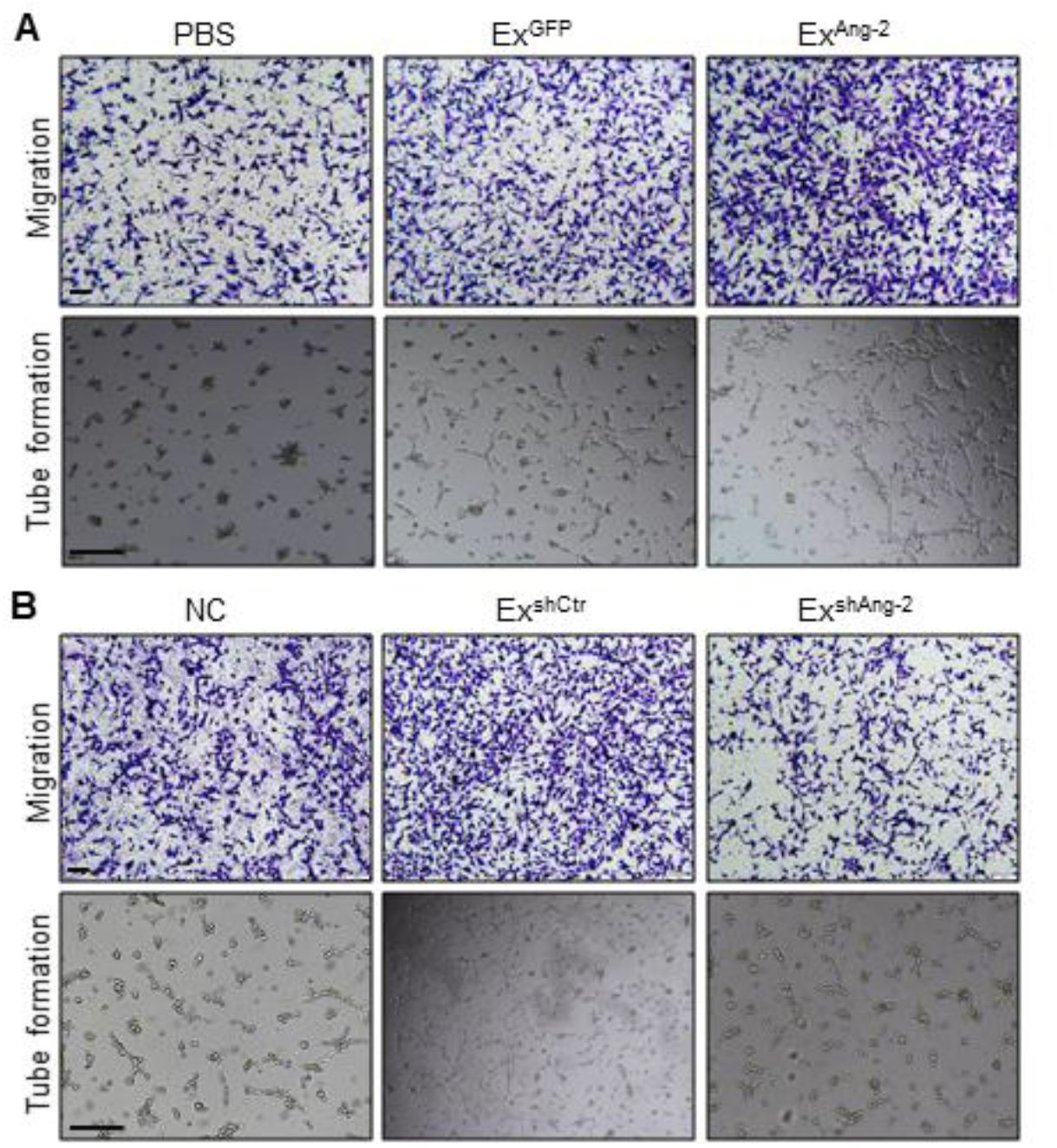
Representative images of cell migration and tube formation in Ex^GFP^, Ex^Ang-2^, Ex^shCtr^, and Ex^shAng-2^ treated HDLECs. A Ex^Ang-2^ further improved HDLEC migration and tube formation in comparison with Ex^GFP^, which has a modest effect. B Ex^shAng-2^ diminished the promotion of Ex^shCtr^ on HDLEC migration and tube formation. Original magnification for HDLEC migration 100×; Original magnification for HDLEC tube formation 40×. Scale bars, 100 μm.

**Figure EV5.**
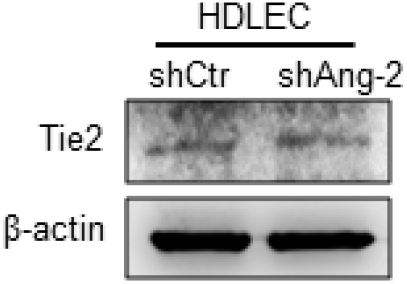
Western blot analysis of Tie2 protein in shCtr-, and shAng-2 transfected HDLECs. Tie2 expression in HDLECs was not interfered by shAng-2 in comparison with shCtr.

